# Extreme ecological specialization in a rainforest mammal, the Bornean tufted ground squirrel, *Rheithrosciurus macrotis*

**DOI:** 10.1101/2020.08.03.233999

**Authors:** Andrew J. Marshall, Erik Meijaard, Mark Leighton

## Abstract

The endemic Bornean tufted ground squirrel, *Rheithrosciurus macrotis*, has attracted great interest among biologists and the public recently. Nevertheless, we lack information on the most basic aspects of its biology. Here we present the first empirical data on the feeding ecology of tufted ground squirrels, and use data from 81 sympatric mammalian and avian vertebrates to place it within a broad comparative context. *R. macrotis* is a dedicated seed predator and shows much more extreme ecological specialization than any other vertebrate, feeding on a far smaller subset of available plant foods and demonstrating a greater reliance on a single plant species– *Canarium decumanum–*than any other vertebrate taxon. Our results suggest that *R. macrotis* plays an important, previously unknown role in the ecology of Bornean lowland forests, and highlight how much we have yet to learn about the fauna inhabiting some of the most diverse, and most severely threatened, ecosystems on the planet.

## Introduction

The Bornean tufted ground squirrel, *Rheithrosciurus macrotis*, has been one of the most talked-about squirrels in recent years. This began with its characterization in the journal *Science* as the ‘vampire squirrel’ [1], which followed an account of local folklore that alleged these squirrels kill deer [2]. The moniker ‘vampire squirrel’ spread widely on social media across the globe. Interest in the squirrel spiked again in 2015 with the release of the first video recordings of these squirrels in the wild at Gunung Palung National Park in West Kalimantan [3]. Despite this global attention, *R. macrotis* remains a very poorly known and largely unstudied species. What we do know is that the species is unusual in many respects. Firstly, phylogenetically it is the only species in SE Asia related to the Sciurini tribe, a large group of Holarctic and South American squirrel species. How *R. macrotis* colonized Borneo remains unclear because there are no known fossils that link it with the other Sciurini from which it separated some 8.6 million years ago [4–6]. *R. macrotis* also stands out because of its unusual incisors in both the upper and lower jaw, which bear a number of deeply carved ridges (~ 10) so that the incisors’ cutting edge is saw-shaped, an arrangement apparently not recorded among other mammals [7]. Its species name likely links to this feature, with the Greek *ρ∊ίθρο* meaning gutter or groove. In addition, comparative morphometric analyses of squirrel mandibles show that *R. macrotis* is a dramatic outlier compared to other squirrels, particularly in its short, robust mandibles with short, wide articular processes [8]. Finally, *R. macrotis* appears to have the largest tail relative to body size of all mammal species, a possible anti-predator adaptation [2].

Although *R. macrotis* is a biogeographic enigma and morphologically unique, little of its basic ecology is known. To our knowledge there has not been any systematic field study of the species’ ecology, although it has been recorded on camera traps at several sites in Borneo (e.g., [9,10]). The large size and unusual shape of *R. macrotis* skulls [8, 11] coupled with their extremely stout incisors and powerful masseter muscles [12, 13] suggest the species is adapted to feeding on extremely hard seeds, but information on the species’ feeding ecology is lacking. Here we present results of a long-term, comparative study of the feeding ecology of an intact community of Bornean rainforest vertebrates to describe the diet of *R. macrotis* and place the degree of its ecological specialization in comparative context. Based on anecdotal published characterizations [14, 15], we hypothesized that *R. macrotis* diets would be dominated by fruit of plants containing hard seeds. In addition, we hypothesized that the extreme morphological specialization of tufted ground squirrels would be reflected in comparatively extreme ecological specialization on a limited subset of available plants. We found that *R. macrotis* is a dedicated seed predator and shows much more extreme ecological specialization than any other vertebrate taxon in our comparative sample.

## Materials and Methods

We analyzed long-term feeding data gathered at the Cabang Panti Research Station (CPRS) in Gunung Palung National Park, West Kalimantan, Indonesia (1°13’ S, 110°70’ E, [16]). Gunung Palung National Park is a formally protected area and all required permits and approvals were secured for the duration of the study from the Indonesian Institute of Sciences (LIPI), the Directorate General for Nature Conservation (PHKA, now KSDAE), and the Gunung Palung National Park Unit (UTNGP, now BTNGP). The study site contains seven tropical forest types, classified based on soil type, drainage, altitude, and parent rock: peat swamp, freshwater swamp, alluvial bench, lowland sandstone, lowland granite, upland granite, and montane [17, 18]. Floristic composition, plant productivity, and mammal densities differ substantially across the habitat gradient [16, 19–21]. During the period of data collection, the site was largely unaffected by hunting or illegal logging, suggesting that the vertebrate and plant communities at the site are characteristic of Bornean lowland forests over recent ecological history.

The feeding data used to describe the diets of *R. macrotis* and compare them to other resident vertebrates were collected by ML and colleagues between March 1985 and March 1992. Trained observers recorded all instances of feeding by vertebrates along standardized census routes through the forest, and opportunistically during other fieldwork. Here we analyze only observations of feeding by vertebrates on fruit or fruit parts (e.g., pulp, seeds), and restrict analyses to independent feeding observations to avoid pseudo-replication (n= 4247, see [18] for a full description of methods). The data set comprises feeding observations for 82 mammalian and avian taxa. Primates (42% of observations) and Rodentia (19%) were the most well-sampled mammalian orders, with additional observations of Artiodactyla (5%), Carnivora (1,6%), and Chiroptera (1.5%). Bucerotiformes (14%), Passeriformes (7 %), and Piciformes (6%) were the best-sampled avian taxa; with additional observations of Columbiformes (1.9%), Galliformes (1.4%), Psittaciformes (0.6%), and Trogoniformes (0.2%).

We compared the dietary diversity of *R. macrotis* with sympatric vertebrates by fitting quadratic plant genus accumulation curves for all vertebrate taxa in our dataset and comparing the species-specific residuals. Vertebrates with large negative residual values were those that fed on a smaller number of plant genera than expected based on the number of feeding observations made (i.e., they specialized on a limited set of plant foods). We used plant genera as the unit for our comparative analyses to account for different levels of taxonomic certainty among plant groups at CPRS [22–24]. To compare the extent of specialization on specific plant genera, we analyzed the dietary composition of all vertebrates for which at least 70 independent feeding observations were available (mean = 274 obs, range 72–549). This resulted in a sample of 12 well-sampled species for comparative analysis. We used the program R 3.3.1 for all analyses and to produce the figures [25]. All code (S1 Code) and data (S2-S6 Data) necessary to replicate our analyses are included as supporting information.

## Results

The field team recorded a total of 79 independent feeding observations for *R. macrotis*, somewhat more observations than for the typical taxon in our dataset (mean= 52, SD = 112, range 1–549 observations/species). These 79 records included feeding on 5 plant taxa; the average number of genera recorded for each vertebrate taxon was 10 (SD = 19, range = 1−88, S2 Data). The most commonly observed taxon in the *R. macrotis* diet was *Canarium decumanum* (n=61 observations), followed by *Mezzetia leptopoda* (n=10), *Elaeocarpus* spp. (n=4), *Dracontomelon* sp. (n= 3), and *Irvingia malayana* (n=1). All *R. macrotis* feeding observations entailed feeding on seeds. Although comparative empirical data on seed hardness are not available, *C. decumanum* and *M. leptopoda* are widely known to be among the hardest seeds found in the Bornean rainforest (AJM, ML personal observations, [26, 27]). Thus, as we hypothesized, the diet of *R. macrotis* is dominated by a small number of plant taxa with extremely hard seeds. Indeed, the plant taxa fed upon by tufted ground squirrels are rarely consumed by any other vertebrates, presumably because their hard seeds are inaccessible to most other taxa [28, S3 Data]. For example, of the 67 independent feeding observations of *C. decumanum* seeds, 59 (88%) were by *Rheithrosciurus*. The only other taxa we observed eating *C. decumanum* seeds were bearded pigs (*Sus barbatus*) and giant squirrels (*Ratufa affinis*), each of which were observed to feed on them 4 times (6%). Both are taxa with powerful jaws known to prey on hard seeds [8, 15, 29]. Of the 15 observations of vertebrate feeding on *M. leptopoda* seeds, 10 (67%) were by *R. macrotis* and the remaining 5 were by bearded pigs (*Sus barbatus*, n=3, 20%) and giant squirrels (*Ratufa affinis.* n=2, 13%, S4 Data).

We also found support for our second hypothesis. *R. macrotis* was extremely ecologically specialized, restricting its feeding to a smaller subset of plant genera than any other vertebrate taxon in our dataset (S2 Data). We observed vertebrate feeding on 159 plant genera. As expected, the number of plant genera recorded in a vertebrate species’ diet increased in a curvilinear fashion with the number of feeding observations recorded (Fig 1, R^2^= 0.97, n= 82 vertebrates, p < 2.2 × 10^−16^). *R. macrotis* exhibited a residual value of −18.2, indicating that it has been recorded feeding on 18 fewer genera than predicted based on the number of feeding observations we recorded. No other taxon exhibits such specialization, with the range of residuals for other species ranging from −8.1 to 11.3. The closest species to *R. macrotis* is the long-tailed parakeet (*Psittacula longicauda*, residual = −8.1), which is also a highly specialized seed predator [29].

**Fig 1.**
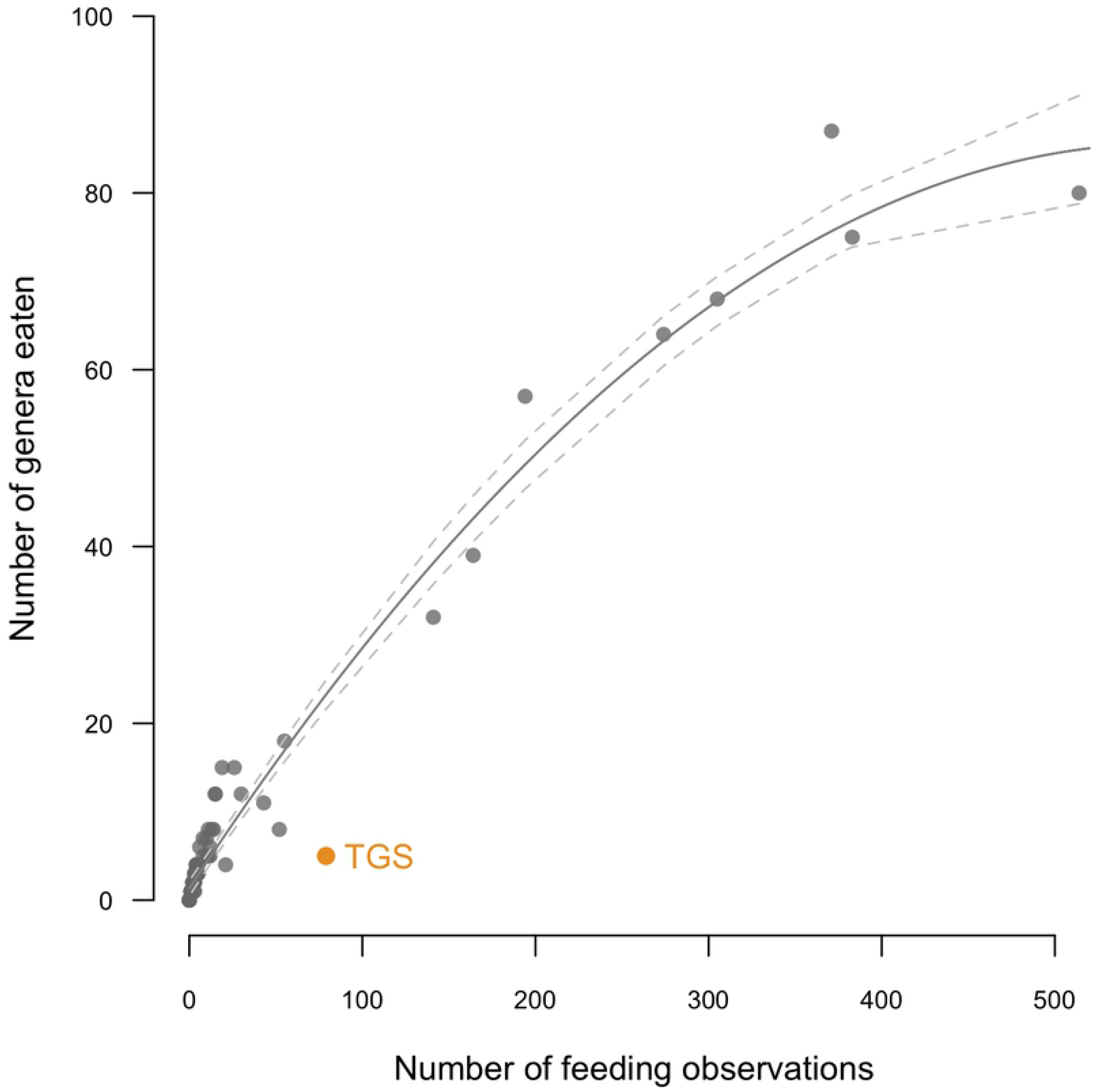
Taxonomic richness of vertebrate frugivore diets. Plot shows the number of plant genera consumed by each vertebrate species plotted against sample size. The plot excludes observations of feeding on the common genus *Ficus.* The curve shows predicted dietary richness (solid line) and the upper and lower 95% confidence intervals (dashed lines). The tufted ground squirrel (TGS, indicated in orange) consumes a much more restricted number of plant genera than expected, and exhibits a much more negative residual than any other taxon.

To further examine the extent of dietary specialization of tufted ground squirrels relative to sympatric vertebrate frugivores, we calculated the importance of each plant taxon in the diets of the twelve most well-sampled vertebrates at CPRS (S5 Data). *C. decumanum* comprised 77% of all *R. macrotis* feeding observations (*C. decumanum* was the only species in the *Canarium* genus consumed by tufted ground squirrels). Only the diverse, keystone plant genus *Ficus* was comparable in its importance in vertebrate diets, comprising more than 80% of the diets of rhinoceros hornbills (*Buceros vigil*) and gold-whiskered barbets (*Megalaima chrysopogon*) and more than 20% of the diets of four other frugivores (Fig 2). No other plant genus comprised more than 22% of the feeding observations of our twelve most well-sampled vertebrate taxa. Interestingly, after *R. macrotis*, the two vertebrates with the largest reliance on a non-fig plant genus were the seed predators bearded pigs (for which *Shorea* seeds comprised 21.9% of feeding observations) and giant squirrels (for which *Lithocarpus* nuts comprised 21.3% of feeding observations). Excluding the extremely specialized tufted ground squirrel, the degree of specialization on non-fig items was not well predicted by overall sample size (β= −0.0001, R^2^= 0.11, n= 11 vertebrates, p = 0.32) nor the number of items recorded in the diet (β= −0.0003, R^2^= 0.02, n= 11 vertebrates, p = 0.68, S6 Data).

**Fig 2:**
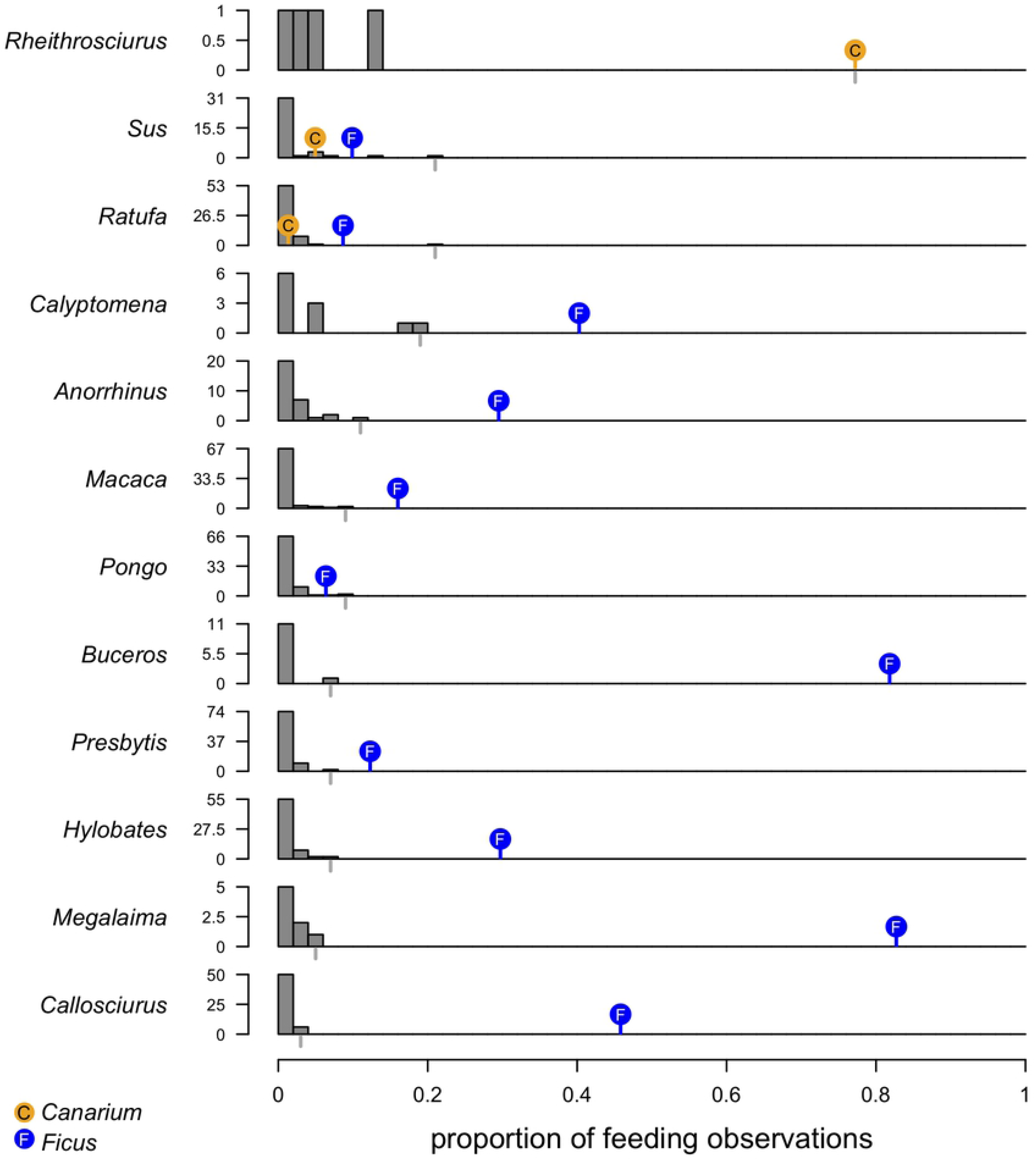
Ecological specialization among well-sampled vertebrates. Histograms of the dietary importance (proportion of total feeding observations) of all plant taxa fed upon by twelve focal vertebrate taxa. Histogram bins are 0.02 in width; y-axis scale depicts the maximum and 1/2 the maximum number of plant items in the tallest bin for each vertebrate. Orange and blue dots indicate the importance of *Canarium* and *Ficus*, respectively. Vertebrate species are listed in decreasing order based on the importance of the most important non-fig plant genus in the diet (indicated by the gray rug line below the x-axis in each panel): *R. macrotis*, *S. barbatus*, *R. affinis*, *Calyptomena viridis*, *Anorrhinus galeritus*, *Macaca fascicularis*, *Pongo pygmaeus*, *Buceros vigil*, *Presbytis rubicunda*, *Hylobates albibarbis*, *Megalaima chrysopogon*, and *Callosciurus prevostii*.

## Discussion

Our systematic study of the feeding ecology of Bornean tufted ground squirrels confirms previous anecdotal descriptions of the species [14, 15]. We provide clear evidence that the species is a seed predator and focuses its feeding on plants bearing extremely hard seeds, especially *C. decumanum* and *M. leptopoda*. Two measures indicate that *R. macrotis* is the most specialized vertebrate taxon in this forest. First, when we controlled for sampling effort, the taxonomic richness of *R. macrotis* diets is far less than that of any other vertebrate frugivore at Cabang Panti. Second, tufted ground squirrels focus on a single plant genus, *Canarium*, far more than any other vertebrate focuses on a single plant genus, with the exception of feeding on the diverse genus *Ficus* (see below). In this context, it is interesting that one of the first descriptions of *R. macrotis* explains that an individual was “[s]hot in the deep jungle during the morning after heavy rainfall, when the animal was looking for fruit under a *Canarium* tree…” ([15]: 125, translated from German). This intense specialization of tufted ground squirrels suggests that they play a crucial role in the ecology of *Canarium* trees–primarily as a seed predator, although likely as an occasional disperser as well in instances where the squirrel buries a seed and fails to return to feed on it (ML personal observation, [15]). In turn, this strong effect on *Canarium* ecology likely has cascading effects on other large canopy tree species that compete with *Canarium*. Specialized seed predators play crucial functions in the ecology of Southeast Asian rainforests [28–30]. Of particular importance are *Psittacula*, *Sus*, and *Ratufa*, which are the taxa in our analysis most similar to *Rheithrosciurus* in their specialization (*Psittacula*) and diet composition (*Sus* and *Ratufa*). This suggests that *R. macrotis* may also play a crucial, heretofore unidentified, role in the ecology of Southeast Asian forests.

*Ficus* is the one plant taxon that exceeds *Canarium* in its importance in the diets of frugivores at the Cabang Panti Research Station (Figure 2). This is not unexpected, both because *Ficus* is a highly diverse plant taxon at this site (56 species in Gunung Palung National Park, [31]) and because it exhibits phenological patterns that make it a keystone species that provides sustenance to a wide range of animals during periods of resource scarcity [32–34]. *Ficus* is widely recognized as a uniquely important plant taxon for tropical frugivores, and it is therefore notable that *Canarium* seeds are of comparable importance to *R. macrotis* in our sample. Unlike *Ficus*, however, *Canarium* is not of major importance to any other vertebrates- suggesting the intensive use of its seeds solely by tufted ground squirrels is a uniquely tightly integrated ecological relationship that deserves further investigation.

While our data present an intriguing story of intense ecological specialization in a rainforest mammal, it is notable that 79 feeding observations constitutes the most complete account of *R. macrotis* ecology ever published. The tufted ground squirrel has attracted an astonishing amount of interest in the scientific and popular press. Erik Stokstad’s news piece in the ‘vampire squirrel’ was the second most read and commented upon article on the website of the journal *Science* during the first week of September 2015–sharing the stage with weighty scientific discussions of “why big societies need big gods” and “how big is the average penis”. *R. macrotis* also possesses several highly unusual features that make it a legitimate subject of serious research attention. In this context, it is truly remarkable how little we know about the species- although our evidence suggests that “assassin squirrel” would be a better moniker than “vampire squirrel”. This highlights our more general ignorance of the biodiversity of Borneo and other parts of the tropics. The vast majority of published research concerns a very limited subset of taxa (e.g., [35, 36]) and with forests in Indonesia and across the tropics being lost at alarming rates [37, 38] we run the risk of losing species before we can collect even the most basic information about their ecology. This is of particular concern for species, such as *R. macrotis*, that are restricted to undisturbed lowland forests and predicted to be highly intolerant of logging and other forms of disturbance [39].

## Acknowledgments

Permission to conduct research at Gunung Palung National Park was kindly granted by the Indonesian Institute of Sciences (LIPI), the Directorate General for Nature Conservation (PHKA, now KSDAE), and the Gunung Palung National Park Office (UTNGP, now BTNGP). We are grateful to Universitas Tanjungpura for counterpart support. We appreciate the assistance and support of the many students, researchers, and field assistants who worked at Cabang Panti Research Station over the past three decades. Special thanks to Emilio Bruna for suggesting the name “assassin squirrel”.

## Supporting information

**S1 Code. All code necessary to replicate the analyses presented in this paper.** All required data are included as supporting information; descriptions of the contents of each data file are provided in this RMarkdown code file.

(Rmd)

**S2 Data. Vertebrate feeding data.** File includes summary feeding information for *R. macrotis* and 81 comparison vertebrate taxa (e.g., taxon, number of independent feeding observations, number of unique plant genera eaten).

(csv)

**S3 Data. Observations of feeding by *R. macrotis.*** File lists all 79 independent feeding observations made by the tufted ground squirrel, classified to plant genus.

(csv)

**S4 Data. Observations of feeding on *Canarium decumanum* and *Mezzetia leptopoda*.** File lists all observations of feeding on two extremely hard-seeded plant species.

(csv)

**S5 Data. Dietary importance of all plant genera consumed by the twelve most well-sampled vertebrate taxa.**

(csv)

**S6 Data. Summary characterization of the diets of the twelve most well-sampled vertebrate taxa.**

(csv)

